# Gauge equivariant convolutional neural networks for diffusion mri

**DOI:** 10.1101/2023.06.09.544263

**Authors:** Uzair Hussain, Ali R. Khan

**Affiliations:** Centre for Functional and Metabolic Mapping, Robarts Research Institute, Western University, London, Canada; Department of Medical Biophysics, Schulich School of Medicine and Dentistry, Western University, London, Canada; Western Institute for Neuroscience, Western University, London, Canada

## Abstract

Diffusion MRI (dMRI) is an imaging technique widely used in neuroimaging research, where the signal carries directional information of underlying neuronal fibres based on the diffusivity of water molecules. One of the shortcomings of dMRI is that numerous images, sampled at gradient directions on a sphere, must be acquired to achieve a reliable angular resolution for model-fitting, which translates to longer scan times, higher costs, and barriers to clinical adoption. In this work we introduce gauge equivariant convolutional neural network (gCNN) layers for dMRI that overcome the challenges associated with the signal being acquired on a sphere with antipodal points identified. This is done by noting that the domain is equivalent to the real projective plane, ℝ*P*^2^, which is a non-euclidean and a non-orientable manifold. This is in stark contrast to a rectangular grid which typical convolutional neural networks (CNNs) are designed for. We apply our method to upsample angular resolution for predicting diffusion tensor imaging (DTI) parameters from just six diffusion gradient directions. The symmetries introduced allow gCNNs the ability to train with fewer subjects as compared to a baseline model that involves only 3D convolutions.

## 1 Introduction

Diffusion weighted imaging (DWI) or diffusion MRI (dMRI) is a type of imaging that conveys microstructural information of underlying tissue, by tracking the diffusion rate of water particles along different directions [1]. Applied to brain imaging, this gives dMRI the unique ability to map the integrity and direction of white matter tracts [2] and also the orientation and dispersion of neurites in grey matter [3]. Diffusion tensor imaging (DTI) is the most commonly used model applied to dMRI datasets, and it aims to predict the diffusion tensor at each location of the image [4]. The eigenvalues of the tensor provide useful rotationally-invariant micro-structural metrics, such as fractional anisotropy (FA) and mean diffusivity (MD). The first eigenvector of the tensor (V1), provides an estimate of the underlying direction of white matter fibres, and can be used to perform tractography for mapping white matter tracts in the brain.

One of the drawbacks of dMRI and analysis techniques like DTI is that many images are required, one for each direction we want measure the diffusion rate along. Although theoretically the diffusion tensor only requires six directions to be fully determined mathematically, in practice more directions are needed to reduce noise, which translates into longer scan times and higher costs. Recently, many machine learning based methods for dMRI have been proposed to address this issue and reduce the noise generated from using fewer directions [5–9]. The earliest of these models, q-space deep learning (q-DL) [8] was able to show that a simple three-layer multi-layer perceptron (MLP) was able to accurately estimate diffusion kurtosis and neurite orientation dispersion and density imaging (NODDI) [3] parameters with fewer directions thereby reducing scan time by twelve-fold. However, this MLP approach did not take advantage of the inherent spatial correlation between voxels in doing so. Convolutional neural networks (CNNs) are a highly-effective means to leverage spatial correlations present in images. For a single scalar 3D image, one can easily employ 3D convolutions, but for dMRI, at each voxel, we have many signals, one for each direction the signal is sampled from in q-space. Thus, at each voxel the signal is realized on a unit sphere, with directions uniformly distributed on the sphere. Further, because of the inherent antipodal symmetry in the diffusion signals, the domain for the signal is actually the set of all unoriented lines from the origin of ℝ^3^, i.e., the real projective plane, ℝ*P*^2^ [10].

Standard CNNs, developed for computer vision tasks, make use of flat, orientable manifolds, and thus are not readily applicable to the non-euclidean and non-orientable nature of the real projective plane. To circumvent these obstacles, one approach taken by authors is to perform convolutions in the spectral space [5]. This approach has the advantage of easily implementing rotational equivariance by employing Wigner matrices acting on spherical harmonics [11], however, a downside is that moving to spectral space introduces an extra level of processing. Another approach is to limit the domain to six predefined directions, then apply regular 3d convolutions while taking the six directions as different channels [6]. Although this method has the advantage of readily using available 3D convolution toolboxes, the directions sampled must be fixed to these six directions, which is rarely the case in practice. Further, this approach does not have the rotational equivariance property. Another promising approach is to use models developed for natural language processing (NLP) like the attention based transformer architecture [7, 12]. However, this requires substantially more subjects for training, which is infeasible in many research or clinical applications. Also, similar to [6], it requires six predefined directions and lacks rotational equivariance.

In this work we apply the theory of gauge equivariant CNNs (gCNNs) [13, 14] to introduce convolutional layers for the real projective plane, ℝℙ^2^, and apply these to the problem of denoising DTI metrics derived from six directions. Using gCNNs allows us to perform convolutions on non-euclidean manifolds, and by modelling ℝ*P*^2^ as the top half of an icosahedron we are able to take convolutions on a non-orientable manifold as well. In addition, we are not limited to a predefined and fixed set of diffusion gradient directions, as our approach can be applied with any set of uniformly distributed directions. With the accompanying open-source implementation, these generalizations should allow the gCNN layers developed here to be applied to other dMRI related problems or architectures where the signal is acquired on a unit sphere with antipodal symmetry. As we will see below, a particularly beneficial side-effect of using gCNNs is a higher learning efficiency brought about by manually rotating and reflecting convolution kernels, as opposed to learning these transformations, and as a result we are able to reach reasonable accuracy levels with significantly fewer training subjects.

## 2 Theory

The theory of gCNNs is developed in [14], which is then elegantly applied to the icosahedron in [13]. For completeness we review some aspects of the theory relevant to this work, and show how it can be applied to the real projective plane, ℝ*P* ^2^, which is the manifold of interest for dMRI.

### 2.1 Gauge transformations

A gauge usually means a choice of some reference point. E.g., in a literal gauge like a speedometer, we calibrate the needle to a zero on the markings, and measure speed against that zero. This idea also applies when viewing coordinate systems as a reference, and then coordinate transformations are a type of gauge transformation. One simple example of a gauge transformation is a rotation of the 2d Cartesian plane by some angle.

More formally, following the notation and definitions of [13], we define a gauge as a position-dependent invertible linear map *w*_*p*_ : ℝ^*d*^ → *T*_*p*_*M* where *T*_*p*_*M* is the tangent space of a d-dimensional manifold *M* at point *p*. If {*e*_*a*_} is the standard Cartesian frame of ℝ^d^ then {*w*_*p*_(*e*_*a*_)} is the frame in *T*_*p*_*M*. A gauge transformation, *g*_*p*_ ∈ *GL*(*d*, ℝ), is a position-dependent mapping where *GL*(*d*, ℝ) is the general linear group (the group of invertible *d × d* matrices). To apply the gauge transformation, one composes *w*_p_ with *g*_p_, i.e., *w*_*p*_ → *w*_*p*_*g*_*p*_. In addition, components of a vector, *v*, in the basis {*w*_*p*_(*e*_*a*_)}, transform as 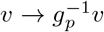, leaving the vector 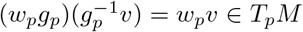 invariant. We can see this invariance in the previous example of the 2D plane and the standard Cartesian basis, now *w*_*p*_ is the identity map, and *g*_*p*_ is a rotation matrix, *R*. Let 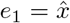 and 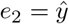. Then we have 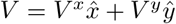 . Under a coordinate rotation we have 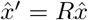, and 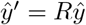, in the new basis 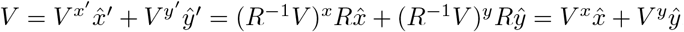.

In general, at a point *p* on *M*, we have other fibres of which *d*-dimensional vectors in *T*_*p*_*M* are a special case. For the purposes of neural networks, a natural object to consider is a feature field *f* with *C* channels. If under a gauge transformation, *f* transforms as 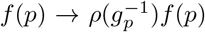, we will call *f* a *ρ*-field where *ρ* : *G* → *GL*(*C*, ℝ) is a representation of *G*. For example, consider a change of coordinate systems in *d*-dimensional Cartesian space by a constant translation, *x*^*i*^ → *x*^*i*^ *+ t*^*i*^. Then a vector *V*^*i*^ transforms as 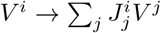, where *J* is the Jacobian matrix.

On the other hand, a complex valued function 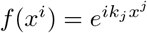, transforms as 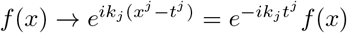. Here the Jacobian matrix is an element of *GL*(*d*, ℝ) but 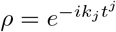 is an element of ℂ (in this particular example we have to extend our field of numbers from real to complex).

To arrive at the definition of gauge equivariant convolutions we need one more ingredient, the exponential map. Let *γ*_*V*_ : ℝ → *M* be a geodesic such that *γ*(0) = *p* and the geodesic tangent is 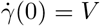 . Then, the exponential map is defined simply by exp_p_ *V* := *γ*_*V*_ (1). This map walks a point along the geodesic starting at *p* with velocity ||*V* || for one unit of time arriving at *q* = exp_*p*_ *V* [15].

### 2.2 Gauge equivariant convolution

Suppose we have a feature map which is a *C*_in_-dimensional *ρ*_in_-field, *f*. We define the following convolution [13]:

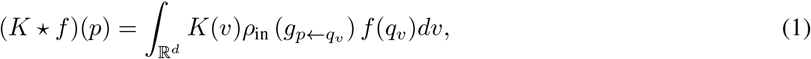

where 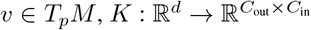 is the kernel, *q*_v_ *= exp*_*p*_*w*_*p*_(*v*) and 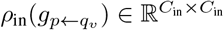 is the transformation that *f*(*q*_*v*_) undergoes as it is transported to *p*. Under a gauge transformation these objects transform as,

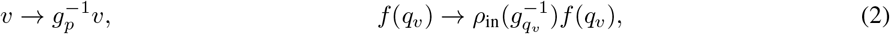

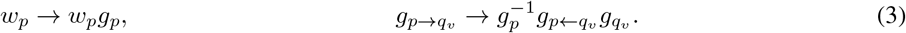

In addition, we require that *K* ⋆ *f* be a *ρ*_out_-field, i.e., it should transform as,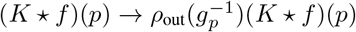 . This is achieved if and only if,

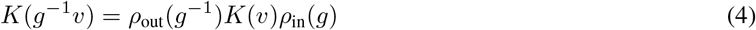

for all *g* ∈ *G*.

### 2.3 Group convolutions of hexagonal filters

In this study we will be using group convolutions with hexagonal filters [16]. We fill first briefly review group convolutions for flat manifolds and then show how it connects with Equation 1. For a group *G*, group convolution is abstractly defined as[14, 17],

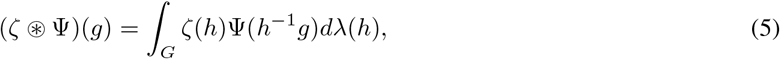

where *g, h* ∈ *G, ζ* : *G* → ℝ is a feature map, Ψ : *G* → ℝ is a filter, and λ is a Haar measure. For hexagonal filters the group *G* is the dihedral group, *D*_6_. As we will see below, since the manifold we are using is the real projective plane, ℝ *P* ^2^, our group, *D*_6_, includes reflections in addition to rotations, giving 12 elements. Writing these elements out, we have,

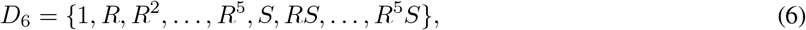

where *R* is a rotation by Δ*θ* = 2π/6. Letting θ_*i*_ = Δθ*i* for *i* ∈ {0, 5}, we then have 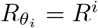. We take *S* to be the reflection along the *θ*_*i*_ = 0 axis; this implies that RS is a reflection along the θ_1_ = Δθ/2 axis, *R*^2^*S* is a reflection along the *θ*_2_ = 2Δ*θ*/2 axis, and so on. To match the notation of the rotations we will denote reflections as 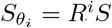, but note that this does not mean that the reflection is along the θ_i_ axis, but rather half of that. Consider the inverses of these operations,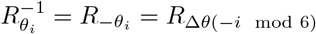. For the inverse of reflections we have 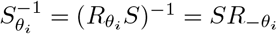. This can be simplified further, noting that 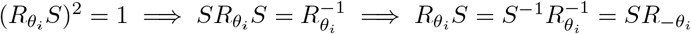. The first equality comes from the fact that if you apply reflection twice it’s the same as identity, then the rest follows. This means that 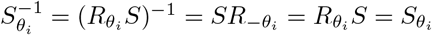.

With *x* = (*x*^1^, *x*^2^) ∈ ℝ^2^, let *I*_*c*_(*x*) be the *c*-th channel of the input and 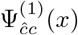 be the hexagonal filter for the first layer [13, 16]. Group operations on the filter are denoted as 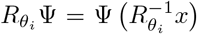 (omitting indices for clarity), and 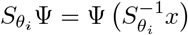. Then the group convolution for the input (scalar) layers is given by [18],

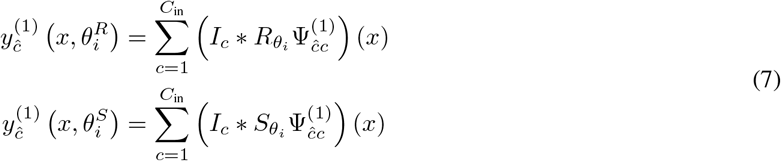

where ∗ denotes a typical spatial convolution, 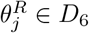 denotes the application of rotation, and 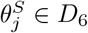 denotes the application of a reflection. After the convolution, we add the bias, *β*_*ĉ*_, and apply a non-linearity *σ*,

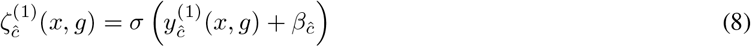

Where 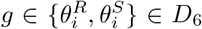. Layers after the first one are called group-convolutional, or “regular” layers, since the input now has both a spatial argument and rotation/reflection argument. This means that the filters will also have rotation/reflection arguments, Ψ^(*l*)^(*x, g*). For these layers, consider first the convolution where we restrict the filter only over rotations,

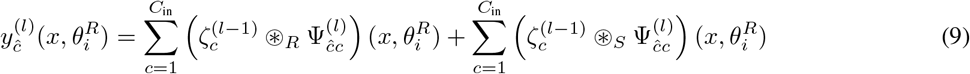

where,

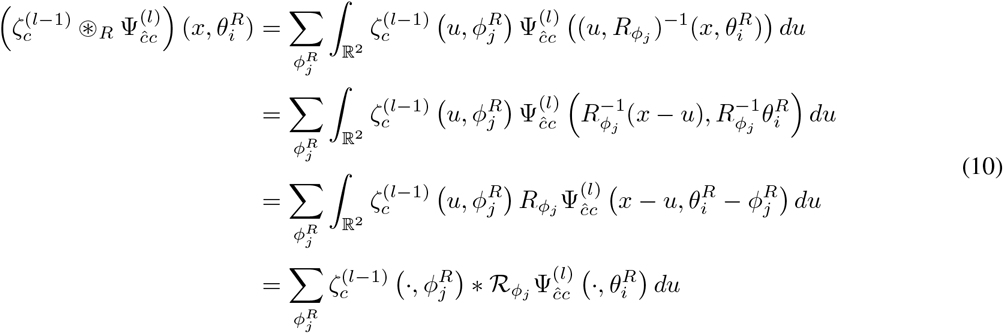

Going from the first line to the second we have used the group multiplication rule, i.e.,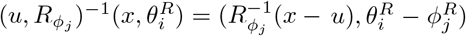 . The third line follows from our notation for rotated filters, and in the fourth line, ∗ denotes spatial convolution and we introduce the notation,

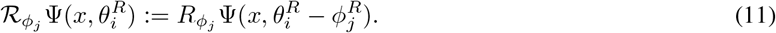

Following similar steps we can show that,

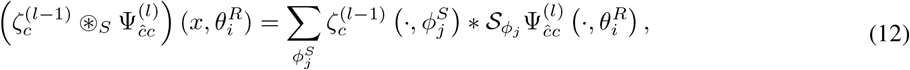

where,

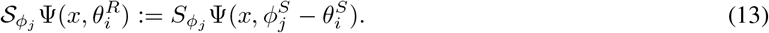

Restricting the filter argument over reflections we get,

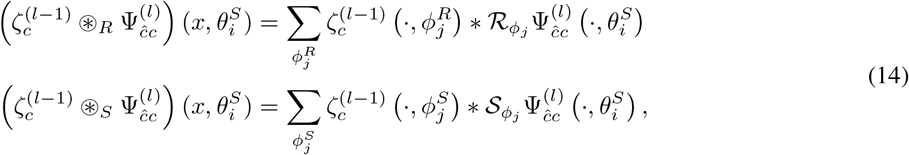

where,

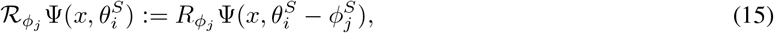

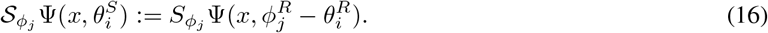

Note that in the Equation 9 we inevitably have two summations, one over the input channels and one over the group elements. To make the connection with the gauge equivariant convolution, Equation 1 we will stack these summations into one channel dimension[13]. Then we have at point, *p*,

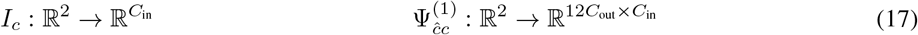

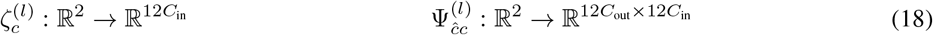

This stacking allows us to express the group convolution in the form of Equation 1 with 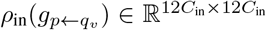. In Figure 1 we see this stack of filters, where the first column shows how a filter, Ψ^(1)^ is expanded for scalar inputs, i.e, rotations/reflections of the the filter. For regular layers, the treatment is different as now we also have an extra dimension *θ*, denoted by different colors in Figure 1. The rows show the action of operators 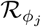 and 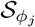 . Notice how *θ*_R_ and *θ*_S_ permute within their blocks with actions of ϕ_R_, but the blocks as a whole also permute with action of ϕ_S_. This is because a rotation followed by a reflection produces a reflection, and two consecutive reflections produces a rotation.

**Figure 1:**
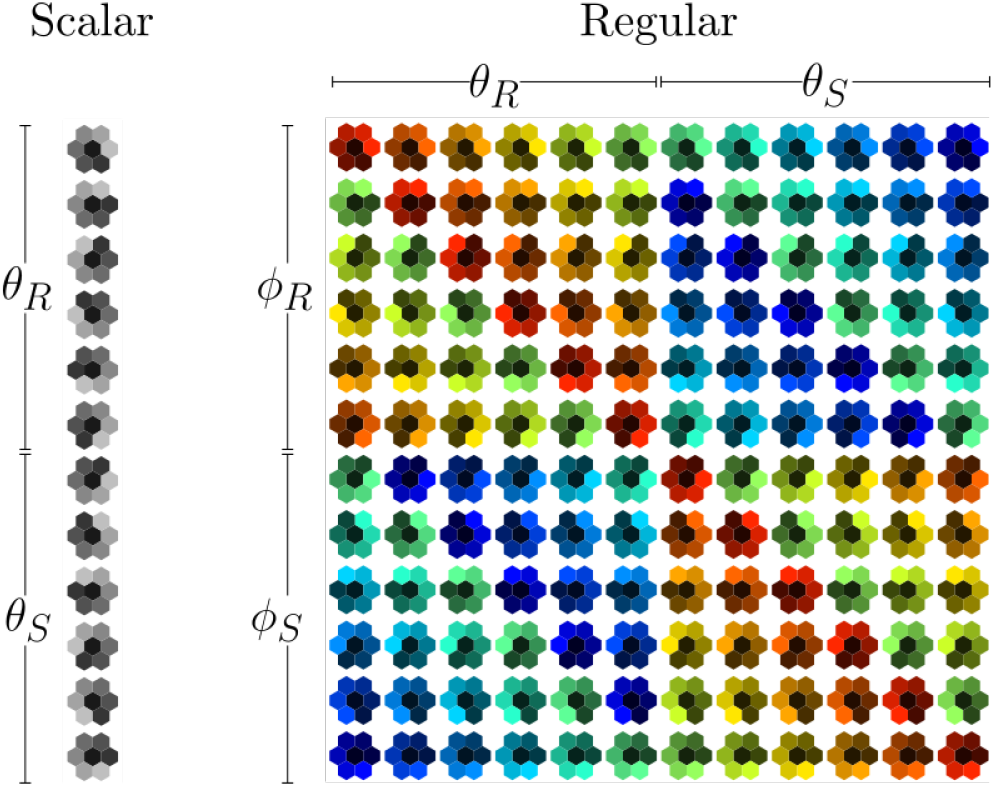
This is an illustration of the group operations on an input layer filter, i.e., scalar layer, and deeper (regular) layers. For the scalar layer we see a simple rotation and reflection of the hexagonal filter. On the other hand, for regular layers, we have an extra dimension, *θ* for previously applied group operations. This is depicted horizontally with the different colors. The vertical axis shows the effect of the operations ℛ_ϕ_ and 𝒮 _ϕ_. Note how a previously applied rotation followed by a reflection produces a reflection, and a previously applied reflection followed by a reflection produces a rotation.

### 2.4 The half icosahedron and padding

In dMRI the signal is assumed to have the following form [19],

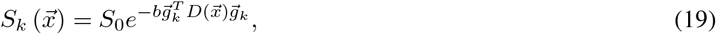

where *S*_0_ is a constant, 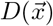 is the diffusion tensor field, 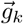 is a unit vector, and *b* is the b-value. In a typical dMRI acquisition there will be many images, *k* ∈ {1, …, *N*_directions_}, such that the unit vectors 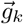 are spread evenly over a sphere. We can see from Equation 19 that the signal *S*_k_ has antipodal symmetry, i.e., it is invariant under the transformation 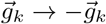. Hence, we assume that the data from dMRI is a function of all unoriented lines from the origin of ℝ^3^, i.e., the real projective plane, ℝ*P*^2^. A topological model of ℝ*P*^2^ can be constructed by a sphere with antipodal sides points identified and then extracting the top hemisphere of such a sphere [10]. This top hemisphere can then be flattened in a manner analogous to the treatment in [13]. This procedure is illustrated in Figure 2.

**Figure 2:**
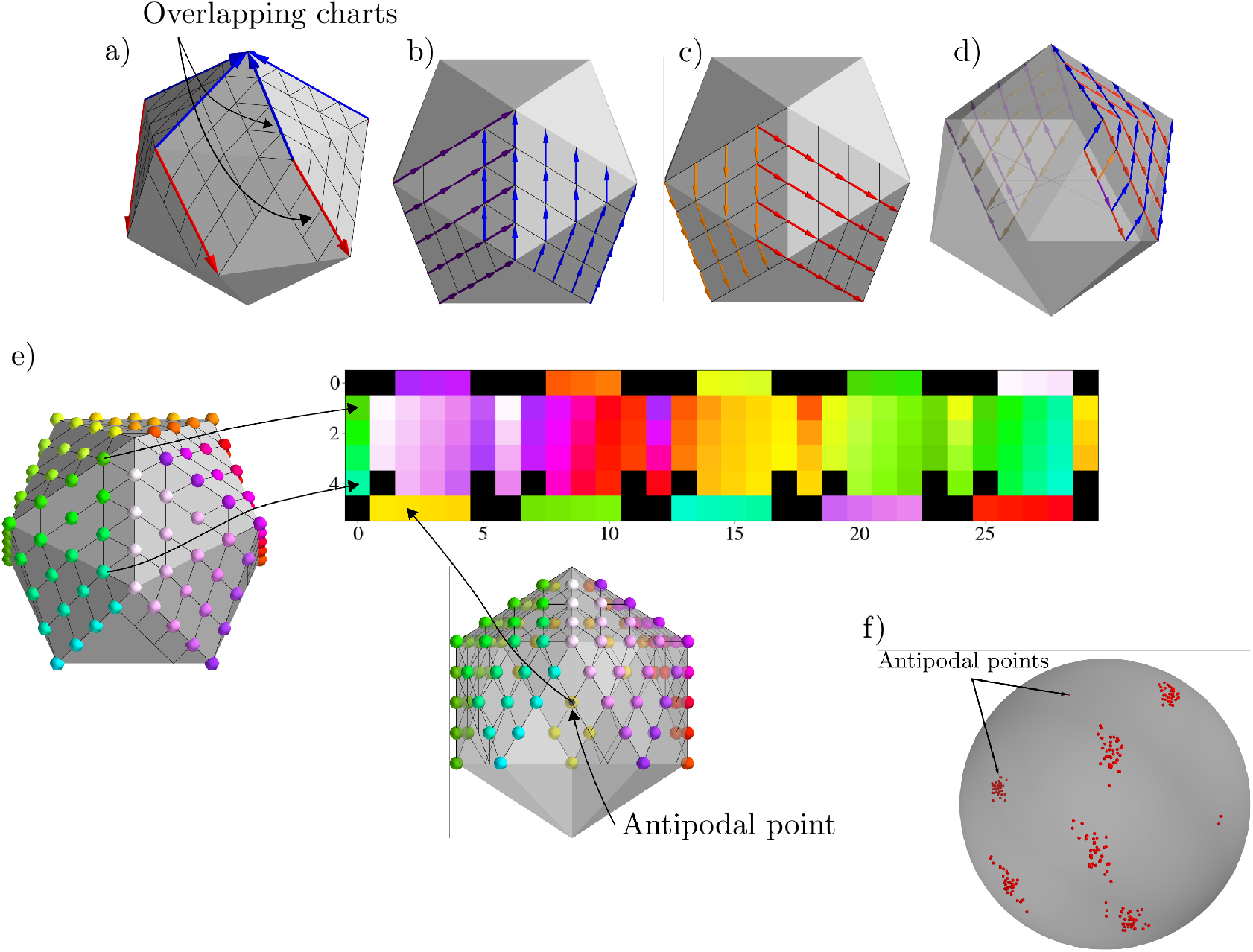
This figure illustrates how a signal and feature maps on the icosahedron are padded, and mapped to a rectangular grid. In a) we see the five square charts with the blue arrow showing the *y*-axis and the red arrow showing the *x*-axis. Also labelled in a) are the regions where the charts overlap, the bottom strip overlaps due to antipodal identification. In b) and c) we see how the unit vectors, blue and red, are related to the unit vectors purple and orange, respectively, of the chart on the left by counter-clockwise rotation of 60°. In d) we see an example of how antipodal identification on the bottom strip leads to a reflection that swaps the *x* and *y* axes. In e) we see how a scalar signal is mapped to the rectangular grid. Note that since there is a strip added on the bottom and left of each 5*×*5 grid, the resulting grid is 6 *×* 30. The padding on these added strips is also illustrated in e). In f) we see the first six directions for the HCP dataset for 40 training subjects, note that there is a small amount of variability in these directions.

Following the procedure of [13], a hexagonal grid on the half icosahedron ℐ can be constructed from series of grid-refinement steps. Let ℋ_0_ be the grid consisting of only the corners of ℐ. We may then take each triangular face and subdivide it into four smaller triangles which introduces three new points on the mid-points of the edges of the original larger triangles. This process can be repeated *r* times to obtain a refined grid, ℋ_*r*_.

The half icosahedron, *I*, has an atlas that consists of five overlapping charts. A chart is an invertible map *φ*_*i*_ : *U*_*i*_ → *V*_*i*_, where *U*_*i*_ ⊂ *ℋ*_*r*_ ⊂ *ℐ* and *V*_*i*_ ⊂ *ℤ*^2^. For each chart we define an exterior 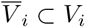, which consists of border pixels, and an interior 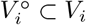, consisting of all pixels that have neighbours in the same chart *i*. To ensure that each pixel (except for the 11 corners) in ℋ_r_ is occupied by a chart, a strip is added added to the left and bottom of each chart as shown in Figure 2. Note in Figure 2a) that some points in the bottom strip are outside the half icosahedron, ℐ, but due to antipodal identification in our topological model they are identified with pixels on ℐ.

For a point *p* ∈ *U*_*i*_ we choose a gauge, 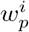, which is illustrated with red (x-axis) and blue (y-axis) in Figure 2a). Let *e*_1_ = (1, 0) and *e*_2_ = (0, 1) be the orthonormal basis vectors for ℤ^2^, then 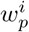 is given by the Jacobian of 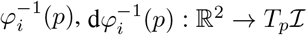. The basis vectors 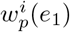 and 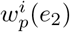 make an angle of 2 · 2π/6 and are aligned with the grid.

Certain pixels, *p* ∈ *U*_i_ ∩ *U*_*j*_, are occupied by multiple charts, for example, in the edges of each chart, either due to sharing an edge or due to antipodal identification. Another example is points on the added strips which are clearly in an overlap of charts, or when some of these points are in multiple charts due to antipodal identification. Note that the basis vectors at such a point *p* may not coincide, i.e, 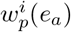 may not be the same as 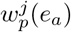 for *a* ∈ {1, 2}. To switch from one chart to another we have to apply a gauge transformation *g*_*ij*_(*p*). In Figure 2b) and c) we see how the *x*-axis (red) and *y*-axis (blue) vectors of a chart compare with the those of a chart on its left. The gauge transformation in this case is a counter clockwise rotation of 60^°^on the left charts vectors to align them on the overlap. This gauge transformation is conveniently an element of the group *D*_6_. Figure 2d) shows how vectors transform as we move to an antipodal point: the x-axis (orange) and y-axis (purple) swap. This gauge transformation, a reflection, is the 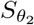 element of the group *D*_6_.

In accordance with these gauge transformations, we can pad the input and the activation maps between convolutions. Figure 2e) shows the padding for a scalar input. There are five square charts, each with dimension 5 *×* 5. Adding the padding strips on the left and bottom, we have dimension 6 *×* 6, and finally stacking them gives dimensions 6 *×* 30. The left strips are padded with neighbouring charts, and the bottom strips are padded with antipodal points. Note that after the first layer, activation maps will have dimensions 6 *×* 30 *×* 12. In this case the padding is similar, but we have to pad from the appropriate θ dimension. For example, when padding the left strip of a particular chart in the θ_*i*_ dimension we take values from the Rθ_*i*_ dimension, when padding the top we take values from the *R*^−1^θ_*i*_, and when padding the bottom and right we take values form the 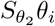 dimension. Note that we pad the right column of each of the five charts after each convolution since the neighbouring column to that is a padding strip for the next chart.

## 3 Methods

### 3.1 Data

We use dMRI data from the Human Connectome Project (HCP) Young Adult 3T study, WU-Minn S1200 release [20]. Informed consent was given by each participant, all methods and experimental protocols had approval from the Washington University Institutional Review Board (IRB) and were in accordance with relevant guidelines and regulations. The dMRI acquisition is multi-shell with b-values of 0, 1000, 2000, and 3000 s/mm^2^ with approximately an equal number of directions on each non-zero b-shell and an echo spacing of 0.78 ms. The resolution is 1.25 mm and the voxels are isotropic. We chose up to 40 random subjects for training and 25 random subjects for testing. We also calculate a brain mask that excludes the cerebrospinal fluid (CSF) using Freesurfer’s `mri_binarize` with `flags --all-wm` and `--gm` (https://surfer.nmr.mgh.harvard.edu) [21–23]. In addition to the dMRI data we also utilize the structural *T*_1_ and *T*_2_ weighted images which are downsampled with cubic spline interpolation to match the resolution of the diffusion images at 1.25mm.

### 3.2 Network

The fundamental network architecture we follow is a residual network [24]. As shown in Figure 3, we have two models; depicted in the grey box is the baseline model which consists of two blocks of 3d convolutions. The first block is denser, followed by another block of 3d convolutions consisting of four layers. Taking the output of the first block of 3d convolutions and passing it instead through four layers of gauge equivariant layers gives the architecture of the second model. Both architectures are trained independently of each other.

**Figure 3:**
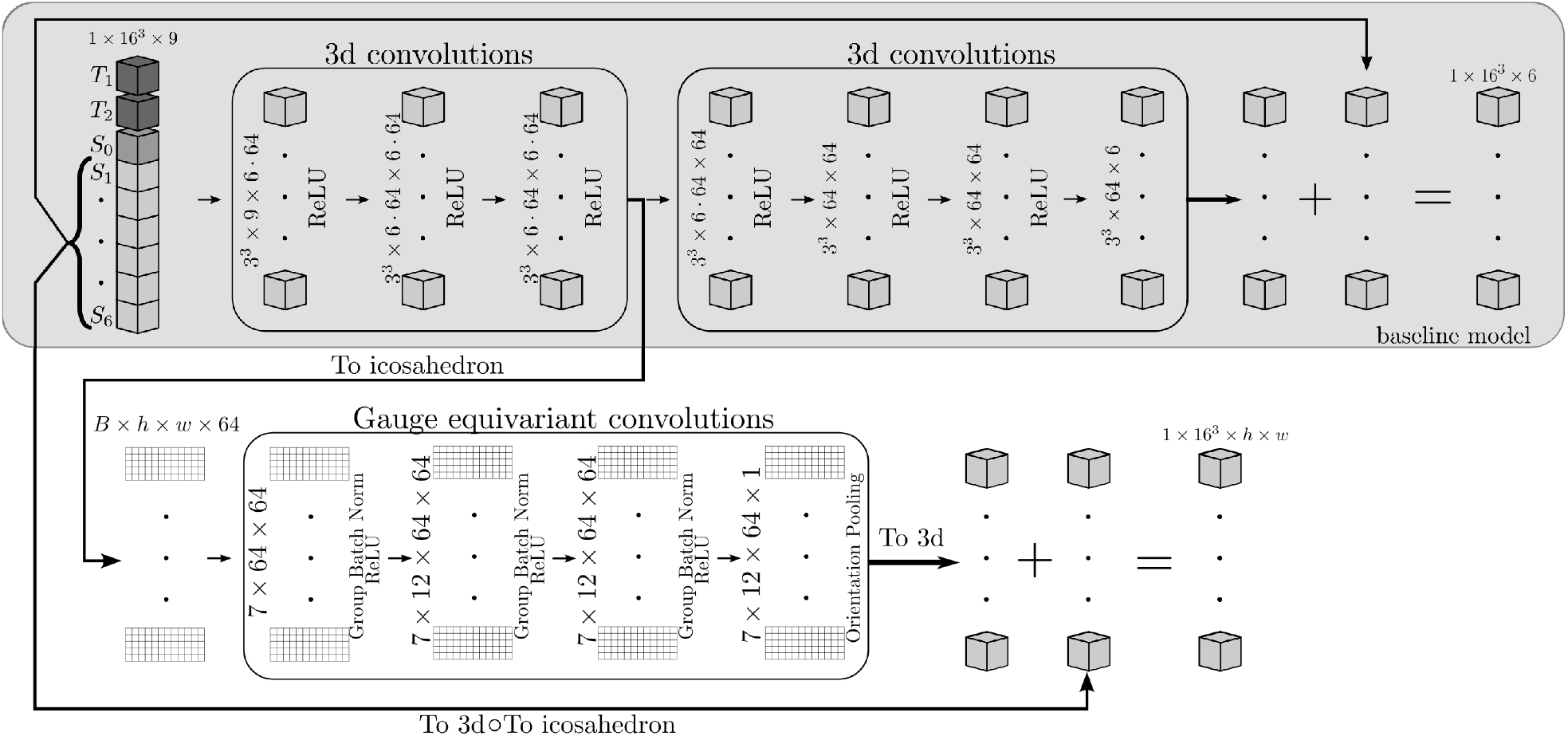
This figure shows the two architectures we used. The input to the network is *T*_1_, *T*_2_, *S*_0_ and *S*_i_ for *i* ∈{1, …, 6} which are the volumes corresponding to six diffusion gradient directions which are simply taken to be the first six in the acquisition to predict an early stop. These nine volumes are broken into 16*×*16*×*16 patches and passed through three layers of 3d convolutions, with each layer containing 6*×*64 filters with a kernel size of 3*×*3*×*3, followed by ReLU activation functions. From here, for the baseline model, we pass the output of the 3D convolutions to another block consisting of four layers of 3D convolutions with 64 filters each. This block has six output channels corresponding to the six input directions, which are then added on. For the gauge equivariant model the output channels of the first block of 3D convolutions are projected onto the top icosahedron mesh which is then flattened. This gives us 64 channels with the 16*×*16*×*16 voxels stacked into the batch dimension. This is then followed by four layers gauge equivariant convolutions with 64 filters each. The first layer has hexagonal filters containing seven weights followed by group convolutional layers with 7*×*12 weights, each layer is followed by group batch norm and a ReLU nonlinearity [13]. At the last layer we perform orientation pooling to collapse the rotation/reflection dimension [18]. Finally we put the batch dimension back into its 3d shape and add the inputs, S_i_ after projection to the icosahedron, giving us volumes with shape 16^3^ *× h × w* where *h* and *w* are the height and width of the flattened icosahedron.

The architecture is similar to the work of [6] with some differences, as opposed to a fixed frame of six directions (or rigid rotations of this frame), the directions are simply taken to be the first six in the acquisition; this is done to mimic an early stop of the scan session. This is possible since the HCP protocol made use of incremental acquisition schemes, which are designed to make sure an aborted scan would result in a reasonably uniform coverage of the sampling domain [25]. *S*_0_ is the first image taken without diffusion gradients applied. To isolate the effects of just the gradient directions, we do not upsample this image and the same *S*_0_ is used during predictions, which is another deviation from [6].

The input is 16 *×* 16 *×* 16 blocks of *T*_1_,*T*_2_, *S*_0_, and six images *S*_1_, …, *S*_6_ corresponding to six gradient directions from the shell with b-value 1,000 s/mm^2^. Given the high memory consumption, we limit our batch size to one. These blocks are then passed through three 3D convolutional layers with `ReLU` non-linearities, filters of size 3 *×* 3 *×* 3 and 6 *×* 64 channels. This gives 64 channels for each of the six gradient directions.

For the gauge equivariant architecture, the output of the first block of 3D convolutions is projected to the icosahedron. This is done by using inverse distance weighting interpolation to sample points on the icosahedron from the six directions for each of the channels. This is followed by averaging antipodal points and then flattening to the top half of the icosahedron as shown in Figure 2. During this flattening, the 16^3^ voxels are moved to the batch dimension with the chosen *h* = 6 and *w* = 30 parameters. This is then passed on to four layers of gauge equivariant convolutions with hexagonal filters [16], i.e., seven weights per channel, 64 channels, followed by group batch normalization layers and `ReLU` non-linearity. In the final gauge equivariant layer, we use an orientation pooling layer to collapse the rotation/reflection direction [17]. Next, the batch dimension is put back into the original 16^3^ shape. Since this is a residual network to get the final output we have to add the inputs. This is done by taking the signal for the six directions, *S*_1_ …, *S*_6_, and flattening it in the same way as the outputs of the 3d convolutions.

We train the models to predict the signal as computed with the tensor model from the fully-sampled data Equation 19, and not the actual acquired fully-sampled signal. We fit the parameters for this equation by using FSL’s `dtifit` https://fsl.fmrib.ox.ac.uk/ [26]. This gives us the diagonalized form of the diffusion tensor *D*, i.e, we get three eigenvectors *V*_1_, *V*_2_ and *V*_3_ and their corresponding eigenvalues λ_1_, λ_2_ and λ_3_. The metrics that are of particular interest are the first eigenvector, *V*_1_, and fractional anisotropy (FA), which can be computed from the eigenvalues and is a measure of how anisotropic the diffusion process is. This approach of fitting to a modelled signal rather than the acquired signal results in more accurate *V*_1_ and FA values after `dtifit` is performed on the predicted signal [6]. For the gauge equivariant model, we generate training labels in the following way, let *X*_flat_ be the projection of the signal at the six directions on to icosahedron followed by flattening to a rectangle as described above. Then we use the dMRI signal for 90 directions with b-value of 1000 s/mm^2^, and perform `dtifit` on it. This allows us to directly compute the signal on the icosahedron mesh using Equation 19, letting this quantity be *Y*_flat_. The final training label is given by *Y*_flat_ − *X*_flat_. For the baseline architecture a similar strategy is used, but the output of the first block of 3D convolutions is followed by four more layers of 3D convolutions with 64 filters each. Here, we also generate the labels using Equation 19 but the signal is sampled along the same six directions as the input, since this is a residual architecture. These input directions are then added to that output at the end.

For the gauge equivariant model training was performed with 5, 10 and 15 subjects, using a learning rate of 1 *×* 10^−4^, reduced by a factor of 0.5 on plateaus, number of epochs was 20 with a batch size of one due to memory constraints, and the loss function as mean squared error. Two V100 GPUs were used, for the maximum 15 subjects training lasted approximately two days. For the baseline model there were two differences; training was performed on 5, 10, 15, 20, 30 and 40 subjects and the number of epochs was 50. The training was significantly faster, less than one day for the maximum of 40 training subjects.

Predictions were performed on 25 test subjects. Once the outputs were computed, for the gauge equivariant model, a single shell dMRI volume was created with a b-value 1000s/mm^2^, with directions taken to be the nodes of the icosahedron mesh. The baseline model simply outputs the same directions as the input. Once we have the volume from both models we perform `dtifit` to obtain FA and *V*_1_. To evaluate the performance of the network in predicting FA and *V*_1_, we compute the mean absolute difference (MAD) between ground truth values and predicted values where the mean is taken over all voxels within the mask that excluded CSF. This is also done for FA and V_1_ values computed from the six direction volume without the enhancement from the network.

## 4 Results and discussion

Figure 4 shows the result of one of the test subjects out of the 25. In a) we have the results from the baseline model and b) the gauge equivariant model. The first column is the input followed by the predictions from each model trained with varying number of training subjects, 20, 30 and 40 for the baseline model in a), and 5, 10 and 15 for the gauge equivariant model in b). The next column shows the ground truth followed by the differences in the last column. The FA difference is an absolute difference whereas for *V*_1_ the difference corresponds to the angle between the vectors. In the *V*_1_ images the direction of the vector is encoded by color as is typical, where red represents left to right, green anterior to posterior and blue superior to inferior.

**Figure 4:**
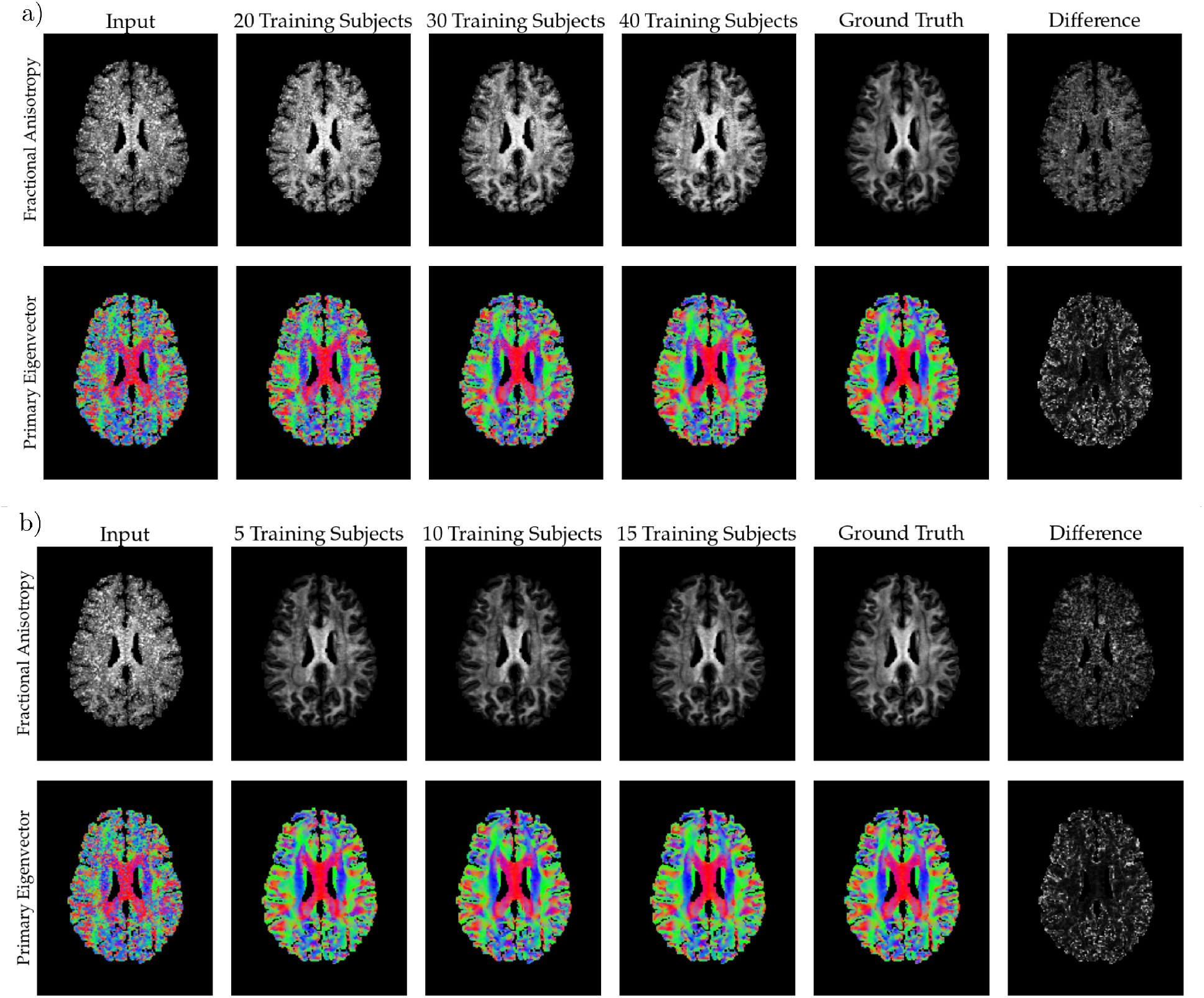
This figure summarizes the results from the networks for one subject. Panel a) shows the results for the baseline model and b) shows the results of the gauge equivariant model. In both a) and b) the rows are FA and *V*_1_, respectively, the first column is the input, the next three columns are the results from increasing training subjects, the fourth column is the ground truth and the last column is the difference between the ground truth and the output from highest number of training subjects, for *V*_1_ this difference is the angle between the vectors. Note that for the gauge equivariant model in b) we show 5, 10, 15 subjects for training. In all panels there is a mask applied that removes the CSF regions. For the primary eigenvector *V*_1_, the colors encode the direction of the vectors, red represents left to right, green anterior to posterior and blue superior to inferior. Below, Figure 5 shows the voxel density plot for the differences as a function of ground truth FA for this subject.

Figure 5 shows the voxel density plots of the subject shown in Figure 4. Here, the CSF voxels are excluded. The *y*-axis in all these plots is the ground truth FA, denoted *FA*_90_. The *x*-axis in a) is Δ*FA* defined as the difference between the ground truth FA, and the FA calculated from the input (first row), the baseline model trained with 40 training subjects (second row) and the gauge equivariant model trained with 15 training subjects (third row). The *x*-axis in b), Δθ, is the angle between the ground truth *V*_1_ and the *V*_1_’s calculated from the input (first row), the baseline model (second row) and the gauge equivariant model (third row).

**Figure 5:**
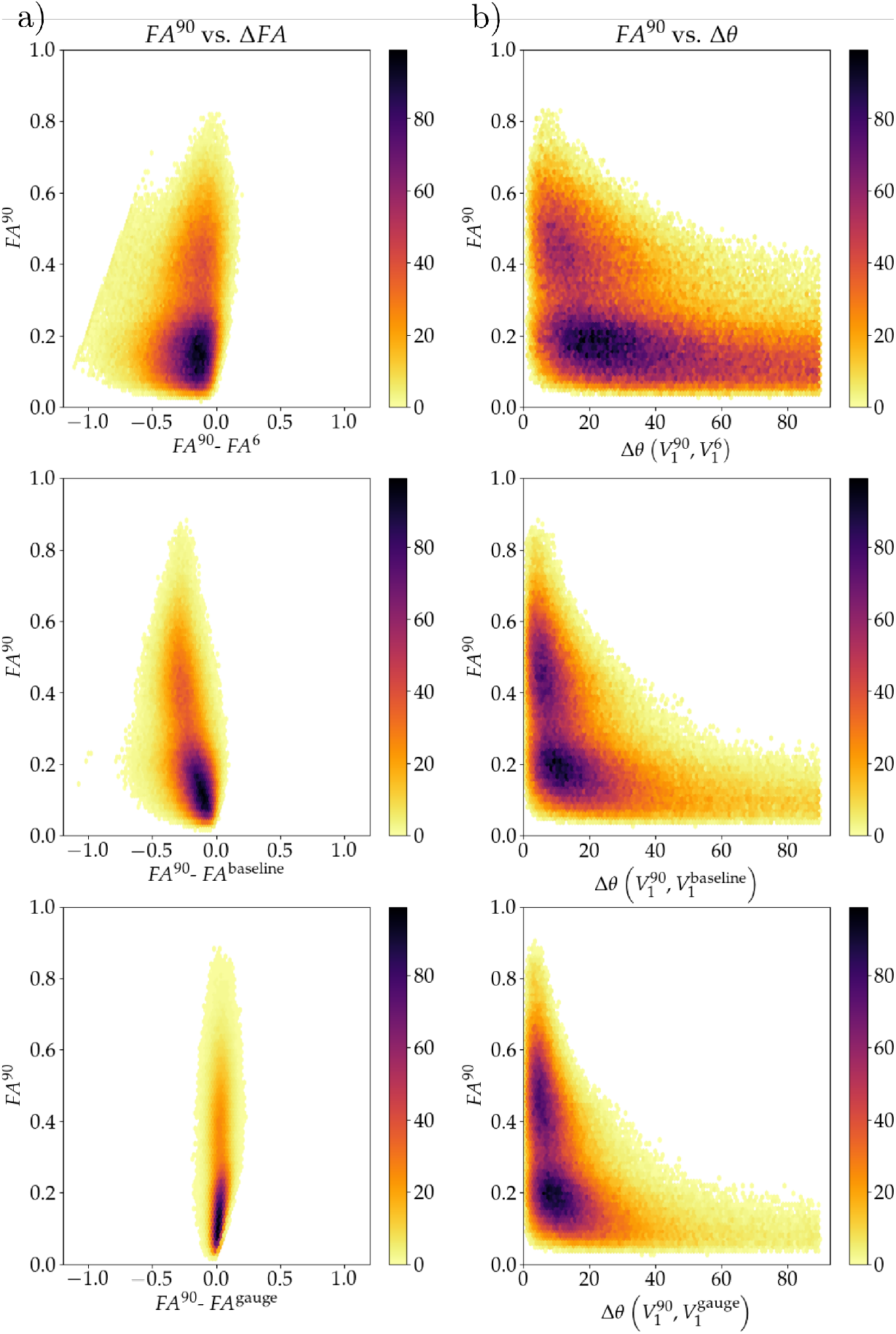
This figure shows the metrics of the model as a density plot over the voxels in the same subject as Figure 4. The three rows in a) and b) correspond to the input, baseline model (40 subjects) and gauge equivariant model (15 subjects), respectively. In all panels the *y*-axis is the ground truth FA, denoted by *FA*^90^. In a) the x-axis is Δ*FA* defined so that Δ*FA* < 0 corresponds to the *FA*^90^ being greater than the input, or model predictions. In b) the *x*-axis is Δθ which is angle between the ground truth *V*_1_ and input, or model predictions.

In Figure 6 we see the effect the size of the training set has on the accuracy metrics for both models. For each of the 25 testing subjects we calculate on the CSF excluding mask the mean over voxels of absolute of Δ*FA* and Δ*θ*, then the mean of these quantities is taken over all subjects. Panels a) and b) show result for Δ*FA* and Δ*θ* respectively, with the orange line for the baseline model and the red line for the gauge equivariant model. Note that due to the relatively high time consumption of the gauge equivariant model, we train up to 15 subjects.

**Figure 6:**
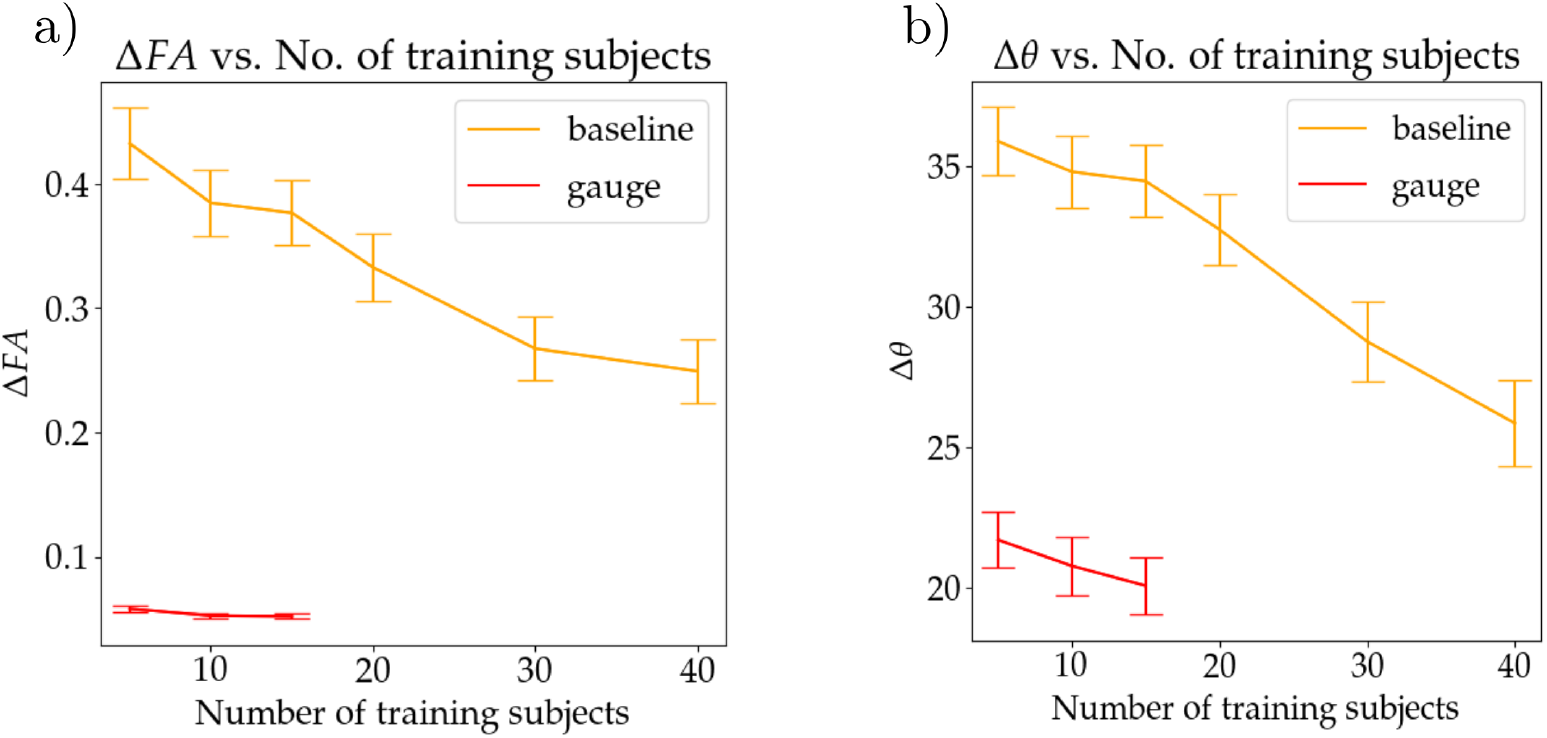
This figure shows the effect of training sample size on the accuracy metrics for both the baseline model (orange) and the gauge equivariant model (red). Panel a) shows the metrics for FA and panel b) for *V*_1_. Each point is calculated by taking the mean absolute difference per subject and then averaging this over all 25 subjects. The error bars show the standard deviation.

Compared to the baseline model that uses just 3D convolutions the gauge equivariant model performs better with fewer training subjects. A particular weakness of the baseline model is that, given the residual architecture, the inputs need to be added to the outputs and hence the number of directions needs to be kept at six, this limits the accuracy of the `dtifit` fitting procedure. Although, as more subjects are added the accuracy of the baseline model improves reasonably well. This limitation is addressed in the gauge equivariant model by projecting the input onto the icosahedron which has many vertices, each vertex corresponding to a volume. Importantly, this projection is done in a manner that is not dependant on the directions that the dMRI signal is acquired on. This coordinate independence on ℝ*P* ^2^ is achieved with an elegant uniting of group convolutions and padding on different coordinate patches on the icosahedron, i.e., the five grids in Figure 2a). As demonstrated in Figure 2f) the first six directions in the HCP acquisition scheme are not completely aligned amongst subjects. In the baseline model there is no straightforward way to address this variability and it is assumed that the six input and output directions correspond to exactly the same directions in all training subjects. In the gauge equivariant model, however, this inter-subject variability is tracked as each subject gets projected on to the icosahedron based on the exact gradient vector. In Figure 6 we see another improvement of the gauge equivariant model which is data efficiency, the model is able to use the training data well and is able to achieve good results with just 15 training subjects. Data efficiency is an expected result of equivariant models [13], and it arises from the architectural requirement for gauge equivariance, where the same convolution kernel is rotated and reflected. These rotated and reflected kernels then don’t have to be learned from more data. This also brings us to the main limitation of the gauge equivariant model, which is its memory and time consumption. Since each kernel has many versions, they have to be saved in memory and convolutions with every version need to be computed. This limitation is what led us to limit training to 15 subjects for the gauge equivariant model (Figure 6). Thus, we see the gauge equivariant approach as a suitable choice in instances where training datasets are small.

## 5 Conclusion

In this work we have laid out the foundations of gauge equivariance that are relevant for dMRI, we achieved this by applying the abstract theory of [13] to ℝ*P* ^2^, where we found that the non-orientable nature of this manifold necessitates reflections of the filter in addition to rotations. We applied our model to the problem of denoising DTI metrics as derived from using only six diffusion gradient directions. We found that the gauge equivariant model performs significatly better than the baseline model that consists of only 3D convolutions. Particularly, since rotated and reflected filters are given as priors and are not learned by the network, the model is data efficient and requires significantly fewer subjects to train as compared to the baseline model. However, this data efficiency comes at a cost of high memory and time consumption, which is a limitation of this approach. We have implemented these layers as an open source package and they can be utilized in other convolutional architectures involving the dMRI signal. In future studies we aim to use these layers for other types of dMRI data like, kurtosis, NODDI, diseases, etc.

## 6 Acknowledgements

Data were provided [in part] by the Human Connectome Project, WU-Minn Consortium (Principal Investigators: David Van Essen and Kamil Ugurbil; 1U54MH091657) funded by the 16 NIH Institutes and Centers that support the NIH Blueprint for Neuroscience Research; and by the McDonnell Center for Systems Neuroscience at Washington University. During the course of this study UH was supported by a postdoctoral fellowship award from the Canadian Open Neuroscience Platform (CONP) and AK was supported by the Canada Research Chairs program #950-231964, NSERC Discovery Grant #6639, and Canada Foundation for Innovation (CFI) John R. Evans Leaders Fund Project #37427, the Canada First Research Excellence Fund, and Brain Canada. This research was enabled in part by the support provided by the Digital Research Alliance of Canada (formerly Compute Canada) (www.computecanada.ca)

## 7 Data and Code Availability

The open source code for this can be found at https://github.com/akhanf/gcnn_dmri. The HCP dataset is openly available at https://www.humanconnectome.org/.

